# Strain-Specific Variation in the Complement Resistome of *Pseudomonas aeruginosa*

**DOI:** 10.1101/2025.01.31.635979

**Authors:** Manon Janet-Maitre, Mylène Robert-Genthon, François Cretin, Sylvie Elsen, Ina Attrée

## Abstract

Bloodstream infections caused by *Pseudomonas aeruginosa* are associated with the highest mortality rates. The complement system, key component of the innate immune response, is mostly responsible for *P. aeruginosa* elimination in human blood. Nevertheless, the plasma sensitivity of *P. aeruginosa* strains varies, ranging from highly sensitive, persistent to fully resistant. Whereas most studies rely on model strains, given the species’ high genomic and phenotypic diversity, the interaction between *P. aeruginosa* and the complement system may be highly complex. In this work, we characterized the plasma resistome of *P. aeruginosa* in three strains displaying variable plasma sensitivity using Tn-seq. The gain-of-function screen performed on the sensitive strain PA14 highlighted numerous bacterial factors impacting plasma resistance, including members of the RetS-LadS/Gac/Rsm regulatory system and outer membrane porins. Tn-seq in plasma-resistant strains CHA and YIK, suggested that each strain utilizes a specific and limited set of proteins to evade complement-dependent killing. Despite this strain-specific response, we identified common strategies across all strains, including the production of exopolysaccharides (EPSs), the presence of surface appendages and modification in the O-specific antigen. We evidence Ssg and Crc as common factors contributing to plasma resistance. However, whereas mutants lacking *ssg* and/or *crc* in plasma had reduced survival, a subpopulation of these mutants persisted over prolonged exposure to human plasma. Overall, this work provides valuable insights into the complex interplay between *P. aeruginosa* and the complement system in the context of bloodstream infections and raises concerns for the development of therapies targeting individual virulence factors.

## Introduction

Bacterial bloodstream infections (BSIs) represent a huge burden for modern healthcare. BSIs caused by *Pseudomonas aeruginosa* exhibit the highest mortality rate (1, 2). This leading nosocomial pathogen causes various primary infections, such as burn wounds, pulmonary and urinary tract infections (3). Using its arsenal of virulence factors, it is able to breach both epithelial and endothelial barriers ((4–7), reviewed in (8)). Once in the bloodstream, *P. aeruginosa* faces the host immune response. In a previous study, we showed the complement system to be the main innate immune component responsible for the elimination of *P. aeruginosa* in the blood (9). The complement system is an enzymatic cascade of more than 30 proteins, which activation results in the formation of the membrane attack complex (MAC) in the outer membrane of the target pathogen. Complement-mediated killing happens through three main steps including complement activation, assembly of the C5b-9 MAC on the outer membrane and penetration of the MAC into the bilayer, eventually leading to bacterial lysis (10, 11). *P. aeruginosa* evolved strategies to evade those three steps. The secretion of proteases including AprA and LasB, cleaving C1, C2 and C3 or the secretion of ecotin, a protease inhibitor, leads to the blockade of complement activation (12–14). *P. aeruginosa* can also recruit host complement inhibitors to its surface, notably through the exposure of the elongation factor Tuf or the dihydrolipoamide dehydrogenase Lpd ((15, 16), reviewed in (8)). Those immune evasion proteins can recruit the Factor H family of complement regulators, resulting in C3b degradation and preventing downstream proteolysis cascade. Finally, *P. aeruginosa* can modify its surface by producing exopolysaccharides (EPS) such as alginate. When acylated, alginates decrease bacterial opsonization by the C3b molecule (17, 18).

Sensitivity of *P. aeruginosa* to the complement system is highly variable. The survival rate of strains in human plasma/serum ranges from full resistance to high sensitivity and persistence and was not found to correlate to the serotype or toxin content (9). Some strains, while mostly sensitive to plasma-dependent killing, persist over a long incubation in plasma (9). Herein, the terms “resistance” and “persistence” will be used as defined in the context of antibiotics with “resistance” attributed to a heritable resistance factor and “persistence” describing a situation where a tolerant subpopulation is able to survive whereas the rest of the population is eliminated (19). A gain-of-function screen identified the first molecular factors involved in *P. aeruginosa* persistence in plasma. However, the screen was performed with the clinical strain IHMA879472 (IHMA87), member of a distant clade of *P. aeruginosa*, recently reclassified as *Pseudomonas paraeruginosa* (20–22).

Several studies documented important genetic and phenotypic diversity of bacterial strains within a *P. aeruginosa* species (22–24), with its pangenome accounting for more than 50,000 genes (22). Given the high intra-species genomic diversity and variability in plasma sensitivity, this work aims at characterizing strategies used by *P. aeruginosa* to evade complement killing in the blood, using genome-wide transposon mutagenesis (Tn-seq) on three strains belonging to different phylogenetic groups (9, 23–25). We found strain-specific sets of factors with only little overlap between strains. However, some common strategies used by *P. aeruginosa* to evade plasma killing could be drawn. We also report that targeting some factors may not eliminate resistant bacteria from human plasma but generate a persistent subpopulation. This work provides insight into a more global comprehension of *P. aeruginosa*-complement system interplay but also raises questions on the targeted antimicrobial or anti-virulence strategies developed on a unique bacterial strain.

## Results

### Choice of strains and global Tn-seq analysis

The three strains used in this work belong to the two major clades of *P. aeruginosa* and have different characteristics in terms of plasma sensitivity, serotype and toxin content (*Figure 1*). The strains CHA, a hyper producer of the EPS alginate, isolated from the lungs of a cystic fibrosis patient (26, 27), and YIK, a recent bacteremia isolate (28), are both resistant to human plasma (∼100% and ∼91% survival, respectively, *Figure 1A*). On the contrary, the laboratory strain PA14 (29) is highly sensitive to plasma and forms a persistent subpopulation, similarly to IHMA87 used in a previous study (*Figure 1A*, (9, 30, 31)). PA14, CHA and YIK harbor the type III secretion system (T3SS) with different toxin content (*Figure 1B*). IHMA87 synthetizes the ExlBA two-partner secretion system instead of the T3SS and is part of the third, the most distal phylogenetic group of *P. aeruginosa* (22, 23), recently reclassified as *P. paraeruginosa* (20).

**Figure 1.**
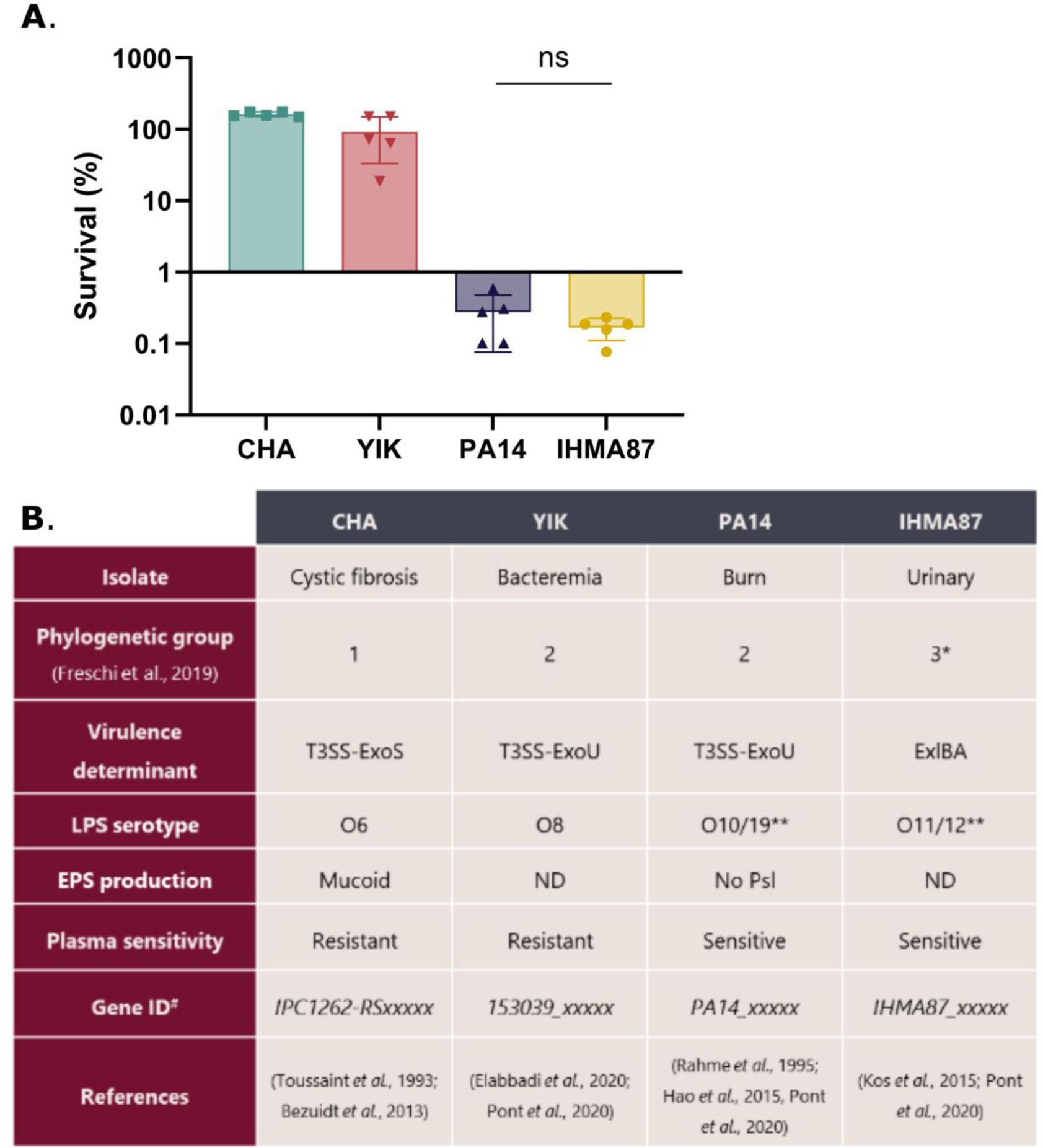
*P. aeruginosa* strains show variable survival in plasma. **A**. Survival after 3h-incubation in human pooled plasma was calculated compared to T0 using colony forming units (CFU) measurements. Data was log-transformed for one-way ANOVA statistical analysis followed by Tukey’s test. All differences in survival between strains were found statistically significant except between IHMA87 and PA14. ns: non-significant. **B**. Characteristics and origin of strains used in this study. ND (not determined). **: reactivity against both serotype antibodies (9, 32). ^#^: Gene ID found in supplemental tables.

We first generated transposon libraries in the three described strains, each containing above 300,000 mutants using the Himar-1 transposon carried by pBTK24 (33). The screen on IHMA87 (21) was done in parallel, as a comparison for PA14. Transposon mutant libraries were challenged for three hours in 90% pooled plasma from healthy donors (output) or heat-inactivated plasma (HIP), unable to kill *P. aeruginosa,* as a control (input). Survivor cells were then recovered in rich medium prior to genomic DNA extraction, bank preparation and Illumina-based sequencing. The set-up of the screen selects for mutants with impaired survival rate, *i.e.* reduced survival for the two plasma-resistant strains (CHA and YIK) or improved survival for PA14. As expected, the overall survival of the pool of PA14 transposon mutants was about 3 log higher than of the parental strain, as already noted for the sensitive strain IHMA87 (*Figure S1*, (21)).

Bioinformatics analysis of the sequencing reads was performed for both coding regions and intergenic regions due to the possible polar effect of the inserted transposon. Depending on the orientation and insertion site of the transposon, which has an outward-directed *P_tac_* promoter, the insertion can either inactivate the target gene or overexpress the neighboring gene(s). The analysis of the Tn-seq data showed that plasma resistance of CHA and YIK relies on few determinants as the screen identified 45 and 37 hits, respectively, using Log_2_(fold change (FC))>1 and adjusted *p-*val (P_adj_)<0.01) as thresholds (*Figure 2 A-D, Tables S1-2*).

**Figure 2.**
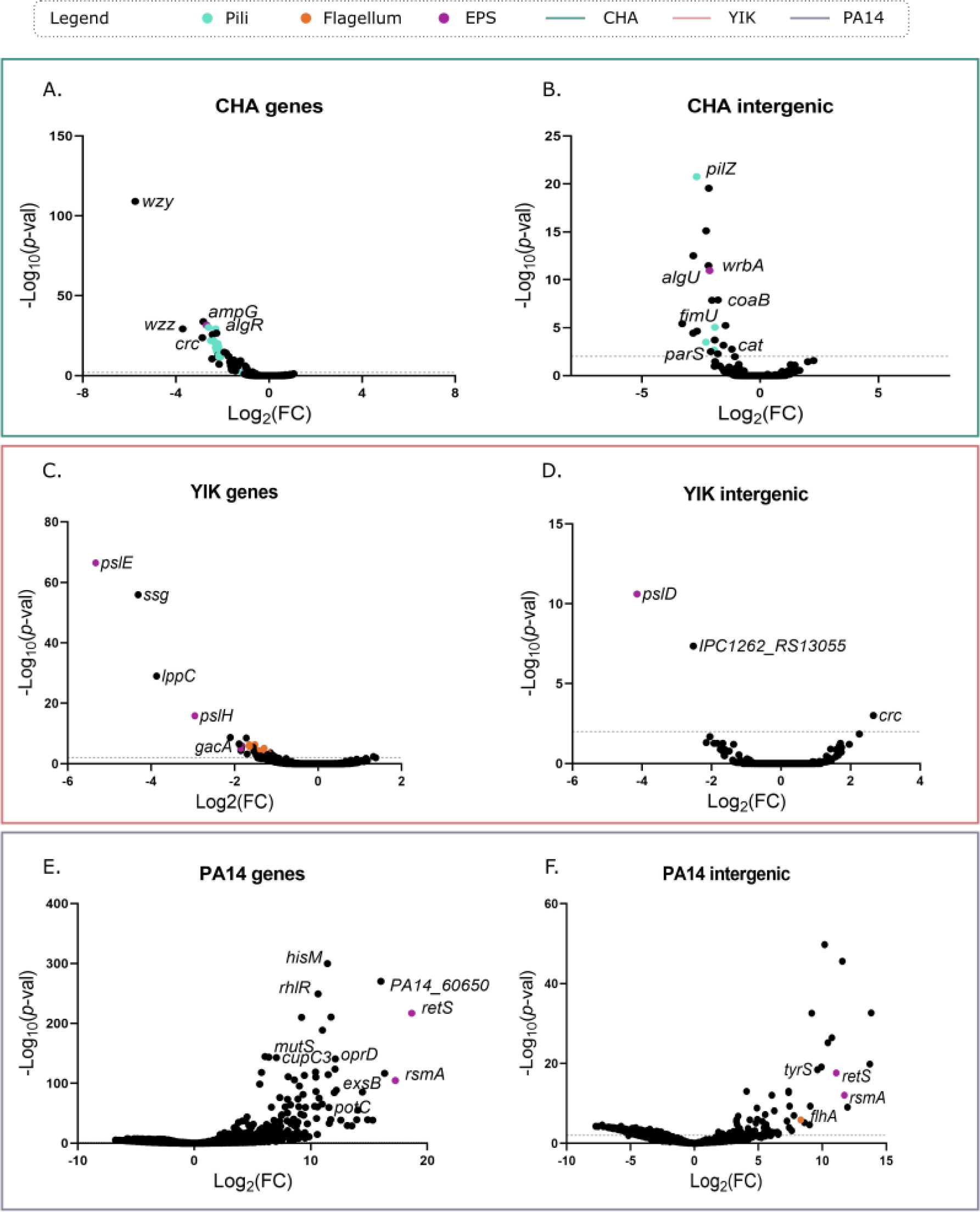
Bioinformatic analysis of Tn-seq data both in genes and intergenic regions. Volcano plot representation of the data obtained for CHA (A-B), YIK (C-D) and PA14 (E-F) both in genes (A, C, E) and intergenic regions (B, D, F). Sequencing data was analyzed using the Galaxy platform (38) and visualized using GraphPad prism 9. Pathways are highlighted in different colors. Dotted line represents a *P_val_* = 0.01.

Conversely, more than 600 hits were found to improve PA14 survival in plasma (682 hits with Log_2_(FC)>1, P_adj_<0.01, *Table S3-4*). As observed on the volcano plot *Figure 2E-F*, few transposon insertions negatively affected PA14 survival in human plasma given the already high sensitivity of the parental PA14 strain to complement-dependent killing. Among them, members of the RetS-LadS/Gac/Rsm regulatory system, involved in *P. aeruginosa* phenotypic switch between planktonic and biofilm lifestyles, had improved plasma resistance including *retS* and *rsmA* with Log_2_(FC) of 18.7 and 17.2, respectively (*Table S3*, (34, 35)). Transposon insertion disrupting the gene of several outer membrane porins, e.g. *oprD*, *oprP*, also drastically improved PA14 resistance to plasma with Log_2_(FC) of 12.1 and 8.0, respectively. The role of OprF, another outer membrane porin as C3 binding acceptor molecule has been previously reported (36). This result suggests that other outer membrane porins may also serve as C3 acceptor. Additionally, disruption of genes involved in metabolisms of nucleotide (pyrimidines: *pyrC, pyrE* and purines: *purH*), aminoacids (e.g. *glyS, hisM*), energy production (*sucC*), biotin biosynthesis (*bioA*) *etc.* led to increased plasma resistance, in line with the data from the strains IHMA87 (21).

Surprisingly, no hit was shared between the two plasma-resistant strains CHA and YIK (*Figure S2*), although the presence and the synteny of most genes were conserved at the genomic level. When comparing the plasma sensitive strain PA14 with the data obtained for IHMA87, 99 common genes showed Log_2_(FC) >1, P_adj_<0.01 (*Figure S2* and (21)). However, except for *retS*, which was one of the top hits for both PA14 and IHMA87 (*Table S3-4*), the other hits had highly variable Log_2_(FC) between the two strains, suggesting that their contribution to protection against the plasma-dependent killing is variable among the strains. The limited redundancy found between the different strains is coherent with what was observed in a similar study in *Klebsiella pneumonia* (37).

Although the individual gene sets were not strictly shared between the strains tested, we examined whether some common strategies were used to evade the complement-mediated killing.

### Contribution of EPS to survival in plasma is strain-dependent

*P. aeruginosa* can produce three types of EPS: Psl, Pel and alginate. Both Psl and alginates were previously described to protect *P. aeruginosa* from opsonization by C3b ((17, 18, 39), reviewed in (8)). Here we confirm EPSs, and particularly Psl, to have a central role in *P. aeruginosa* survival in plasma, but in a strain-dependent manner.

Psl is a neutral exopolysaccharide, which is important in *P. aeruginosa* attachment and maintenance of the biofilm architecture. Although Psl biosynthetic genes are encoded in the same locus, the detailed role of individual genes in Psl synthesis is still unclear (40). Briefly, Psl is thought to be produced by the Psl biosynthesis machinery. PslA is proposed to assemble oligosaccharide repeating units that are then polymerized by PslE. PslG controls the size of the oligosaccharide. Finally, Psl is exported to the bacterial surface through the transporter PslD (*Figure 3A*; (40, 41)). Psl was shown to reduce bacterial opsonization including reduced C3b, C5 and C7 binding at the bacterial surface (39). However, the role of Psl on the MAC-dependent bacterial killing is still under debate. Jones and Wozniak (42) showed a slight beneficial effect of Psl on survival of a *P. aeruginosa* mucoid strains to serum killing, which was not observed by Mishra et al. (39).

**Figure 3.**
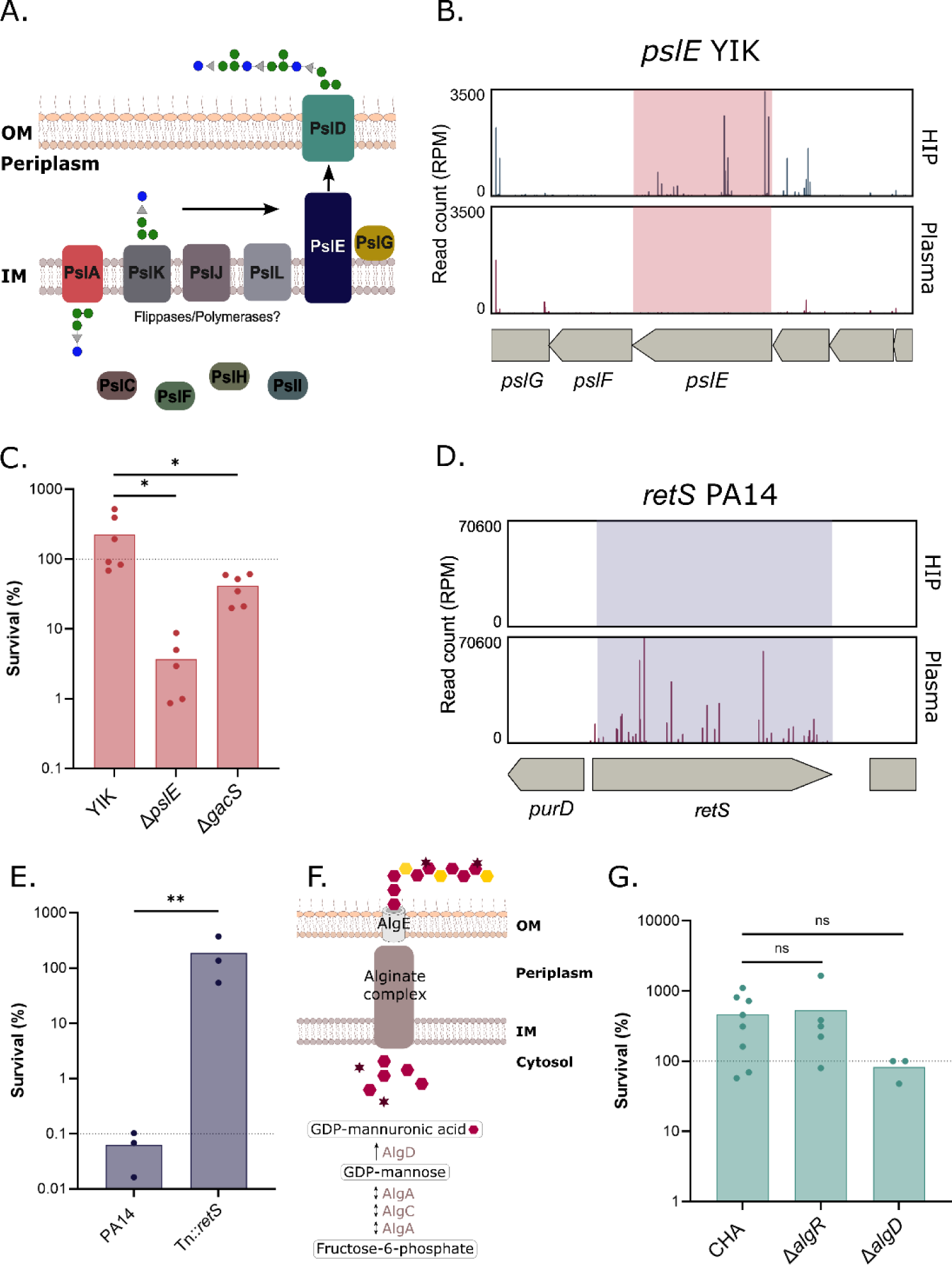
Exopolysaccharides contribute to survival in plasma in a strain-dependent manner. **A.** Scheme of Psl production and export in *P. aeruginosa*. **B.** Tn-seq profile of *pslE* genomic environment in YIK showing number of normalized reads in input (HIP) and output (plasma). **C.** Survival after 3h-incubation in human pooled plasma of YIK, YIKΔ*pslE* and YIKΔ*gacS*. **D.** Tn-seq profile of *retS* genomic environment in PA14, showing number of normalized reads in input (HIP) and output (plasma). **E.** Survival after 3h-incubation in human pooled plasma of PA14 and PA14Tn::*retS*. **F.** Schematic of alginate synthesis and export in *P. aeruginosa*. **G.** Survival after 3h-incubation in human pooled plasma of CHA and CHAΔ*algR* and CHAΔ*algD*. Note that the survival of CHAΔ*algD* is ∼100%. A log-transformation was performed on the dataset and a one-way ANOVA followed by Tukey’s test. Dashed line represents expected survival for the strain shown. ns: non-significant, *: *p_val_*<0.05,**: *p_val_*<0.01.

In the non-mucoid, plasma resistant strain YIK, transposon insertions in three genes within the *psl* locus drastically reduced bacterial survival in plasma (*pslE* – Log_2_(FC)= -5.33, *pslH* – Log_2_(FC)= -2.95, *pslD* – Log_2_(FC)= -1.86, (*Figure 2C-D, Figure 3B, Table S2*). We engineered a *pslE* deletion mutant (Δ*pslE*) in YIK and assessed its survival rate in human plasma. On average, 20 times fewer bacteria were recovered after the exposure of YIKΔ*pslE* to plasma compared to the parental strain, supporting the role of Psl in complement evasion in YIK (*Figure 3C*).

Furthermore, the RetS-LadS/Gac/Rsm regulatory system, which participates in the phenotypic switch between planktonic and biofilm growth and regulates the expression of EPS loci (35, 43), was found a hit in several strains. For example, in YIK, the deletion of *gacS* led to a two-fold decrease in bacterial counts compared to the parental strain when challenged with plasma (*Figure 3C*). Similarly, in the plasma sensitive strain PA14, a transposon insertion mutant in the gene *retS* (PA14::Tn*retS)* displayed a 1000 fold increased survival (Log_2_(FC)=18.66) (*Figure 2EF, 3D-E*), confirming previous data in the strain IHMA87 (Log_2_(FC)=9.11, (21)). The PA14 strain is a natural mutant in LadS and lacks several *psl* genes (44, 45). Therefore, the Tn insertion in *retS* probably activates the synthesis of Pel EPS improving PA14 survival in plasma.

Surprisingly, transposon insertion mutants in either *pel*, *psl* or alginate biosynthetic pathways in hyper-mucoid strain CHA, showed unaltered plasma resistance (*Figure S3*, *Table S1*) Accordingly, a nonmucoid *algD* mutant, in which the last step of alginate biosynthetic pathway is disrupted, (*Figure 3F*) displayed comparable plasma resistance to the parental strains (*Figure 3G*). Previous studies reported that alginates and especially acetylated alginates decreased C3b-mediated bacterial opsonization and therefore increased bacterial survival in clinical strains isolated from cystic fibrosis patients (17, 18), which was not confirmed in our work using the alginate hyper-producer strain CHA. It is worth noting that transposon-insertion mutants in *algU* or *algR* showed decreased abundance following plasma challenge (*Table S1*). Both genes, *algU* encoding the alternative sigma factor and *algR* encoding for a global regulator, are key players in *P. aeruginosa* virulence (46–49). However, the resistance to plasma of an *algR* deletion mutant was found unaffected when tested individually, as opposed to tested in-pool during the Tn-seq experiment (*Figure 3G*). Due to the large size of AlgU and AlgR regulons involved in the response to envelope stress and virulence (47, 49), both proteins may affect CHA plasma resistance in alginate-independent manner. Hence, although alginates can increase plasma resistance in sensitive strains (*e.g.* IHMA87 (21)), they are not crucial EPS for evasion in other strains.

Together, our work shows that the three types of EPS provide potent but strain-dependent protection against the complement system in plasma.

### Surface appendages favor survival in plasma

Bacterial flagella and pili are potent immune activators (50). In this work, we found that the type IV pili (T4P) and the flagellum are surface components providing advantage to strains CHA and YIK in plasma (*Figure 4, Table S1-2*). In the screen with the strain CHA, the transposon mutants in 18 of T4P-related genes had a reduced abundance in plasma compared to the inactivated plasma (*Table S1*). In addition, in the strain YIK, Tn mutants were underrepresented for 12 flagella-related genes (*Table S2*).

**Figure 4.**
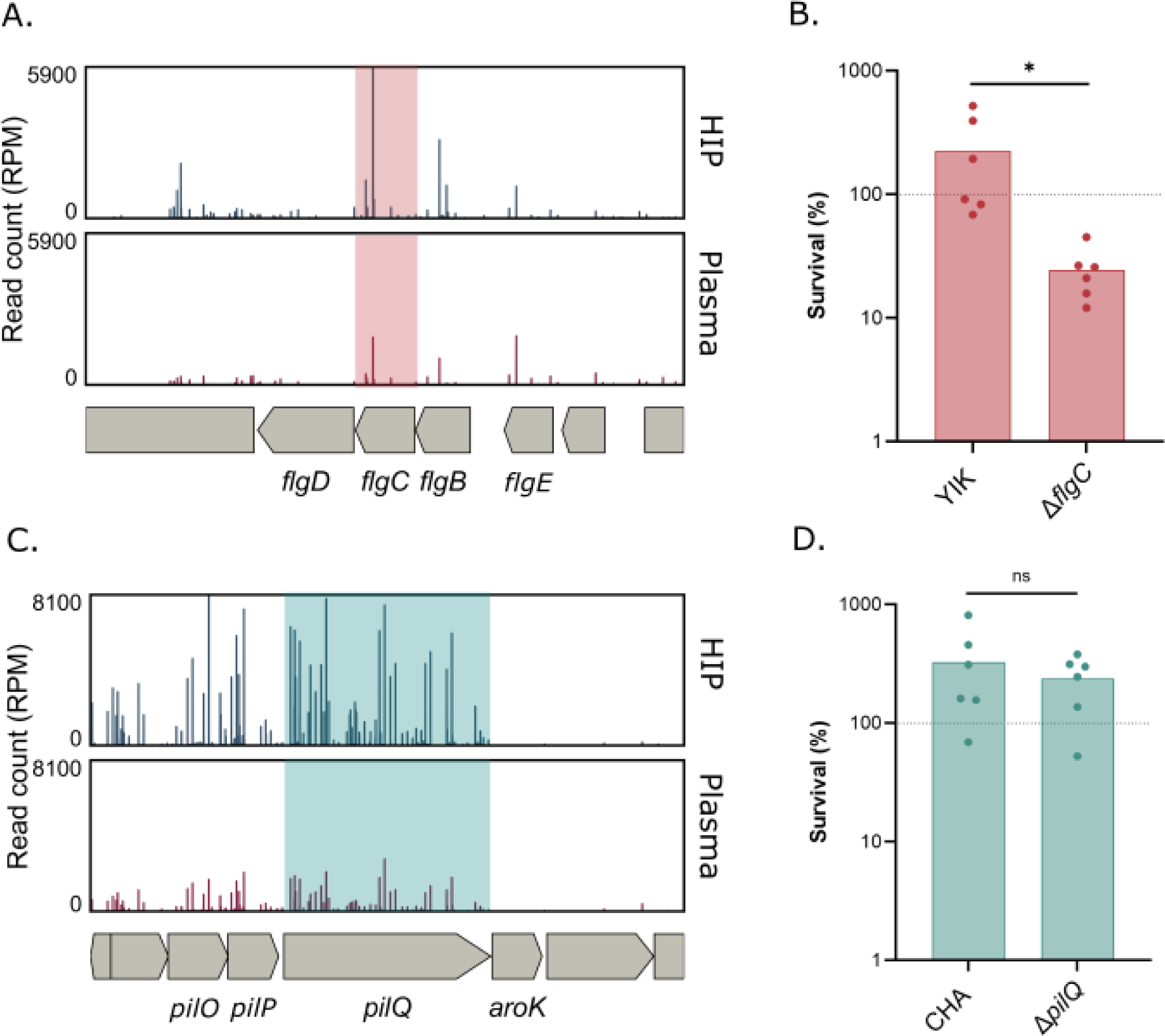
The strain-specific role of surface appendages in survival in plasma. **A.** Tn-seq profile of *flgC* genomic environment in YIK showing number of normalized reads in input (HIP) and output (plasma). **B.** Survival after 3h-incubation in human pooled plasma of YIK and YIKΔ*flgC.* **C.** Tn-seq profile of *pilQ* genomic environment in CHA showing number of normalized reads in input (HIP) and output (plasma). **D.** Survival after 3h-incubation in human pooled plasma of CHA and CHAΔ*pilQ.* Dashed line represents expected survival for the strain shown. The data was log-transformed, and a t-test was performed. ns: non-significant, *: *p_val_*<0.05.

To validate the data, we created a *flgC* deletion mutant in YIK and a *pilQ* mutant in CHA and challenged them individually in human plasma. In accordance with Tn-seq sequencing reads (*Figure 4A*), the deletion in *flgC,* encoding a component of the flagellar basal body (51), resulted in a 4-fold drop in survival of YIK (*Figure 4B*). Surprisingly, the deletion in CHA of a gene essential for pili synthesis, *pilQ* (52, 53), did not confirm the Tn-seq data (*Fig 4C and D*). However, considering the Log_2_(FC) and *p*-values of a number of T4P-related hits in Tn-seq datasets, it is unlikely that pili genes were false-positives. One explanation could be that the identified transposon mutants had a disadvantage in the pooled Tn-seq screen compared to the challenge of an individual deletion mutant.

Several studies point toward a protective role of flagella and pili against serum-mediated killing. In *Streptococcus pyogenes*, pili were shown to bind haptoglobin, an immunosuppressive serum protein, which sequestration provided a survival advantage in serum (54). Tad pili, one type of T4P in *Vibrio vulnificus,* protect bacteria from complement-mediated killing in serum (55). In addition to pili, flagellar components were also found to be required for serum resistance, such as FliG in *E. coli* (56).

Altogether, motility appendages participate in bacterial resistance to human plasma in YIK, by potentially binding serum components (opsonin or other inhibitors) in turn decreasing complement-mediated lysis of *P. aeruginosa*.

### Ssg and Crc mutants persist in plasma

Previous studies showed that MAC insertion in the outer membrane is impacted by the length of O-specific antigens (OSA - (57)). *P. aeruginosa* LPS is synthetized in two forms, either bound with a D-Rhamnose homopolymer known as Common Polysaccharide Antigen (CPA) or OSA, a heteropolymer of sugar repeats with repeat composition determined by the *wbp*_OSA_ locus (58, 59).

The length of newly generated OSA is regulated by Wzz1 and 2, before its attachment to the nascent lipid A (57, 60, 61). In the mucoid and plasma-resistant strain CHA, transposon insertions in *wzy* and *wzz* represented the two best hits and led to elimination of mutants from plasma (Log_2_(FC) = -5.75 and Log_2_(FC) = -3.71, respectively) (*Table S1, Figure 5A-B*), which was confirmed by the complete elimination of clean deletion mutants of either *wzy* or *wzz* in CHA (*Figure 5C*). Similarly, a transposon insertion in the promoter of *wzz,* probably leading to the overexpression of the *wzz* gene, was enriched in plasma in the sensitive strain IHMA87 (21). Although *wzz* was not a significant hit in the PA14 Tn-seq dataset, the zoomed Tn-seq profile suggests a similar role of *wzz* in PA14 plasma resistance (*Figure S4*). Altogether, our data are in agreement with the literature.

**Figure 5.**
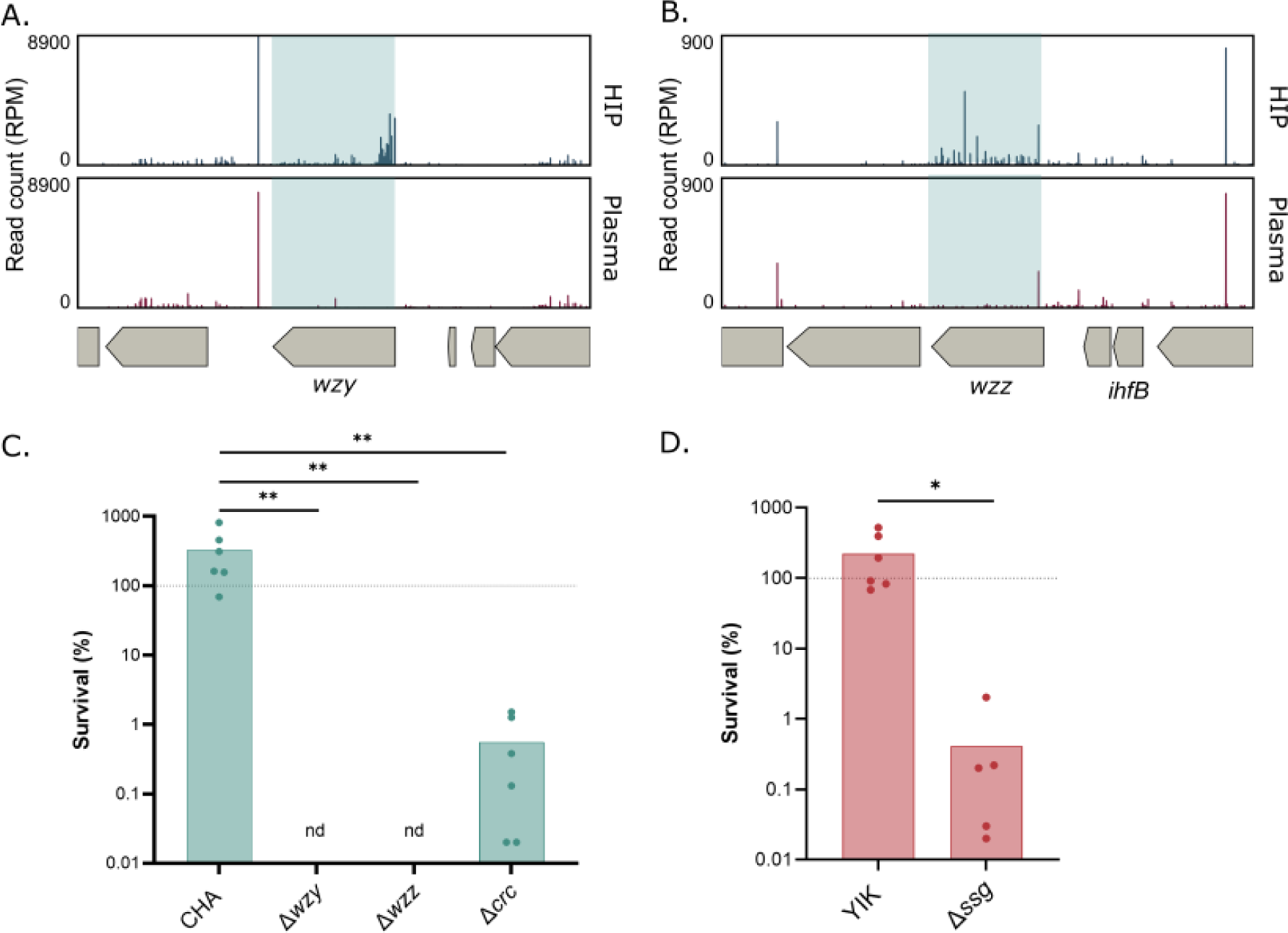
LPS is crucial for survival in plasma. Tn-seq profile of *wzy* (**A.**) and *wzz* (**B.**) genomic environment in CHA showing number of normalized reads in input (HIP) and output (plasma). **C.** Survival after 3h-incubation in human pooled plasma of CHA, CHAΔ*wzy,* CHAΔ*wzz and* CHAΔ*crc.* nd: non detected. Data was log-transformed, and a one-way ANOVA and Tukey’s test were performed. **D.** Survival after 3h-incubation in human pooled plasma of YIK and YIKΔ*ssg.* Data was log-transformed, and a t-test was performed. *: *p_val_*<0.05.**: *p_val_* <0.01. Dashed line represents expected survival for the strain shown.

In addition to *wzz* and *wzy*, two other genes with potential role in LPS synthesis were identified in the screen: *ssg* and *crc*. The inactivation of *ssg*, a conserved gene encoding a putative glycosyltransferase, in the strain YIK, led to a drastic survival drop (Log_2_(FC)= -4.31, *Figure 5D, Table S2*), whereas *crc* with a Log_2_(FC)= -2.86 was the second top hit in CHA (*Figure 5C, Table S1*).

A deletion *crc* mutant tested individually showed a drop in plasma resistance, reaching less than 1% survival after a 3h-challenge compared to the fully resistant parental strain. In the Tn-seq data, *crc* was not found as a significant hit in the YIK strain, nor was *ssg* in CHA. Therefore, we reexamined those genes in the original Tn-seq datasets. Even though the data was not significant due to less insertion mutants in the input library, a similar trend could be observed in both strains (*Figure 6A-B* and *Figure S5A-B*).

**Figure 6.**
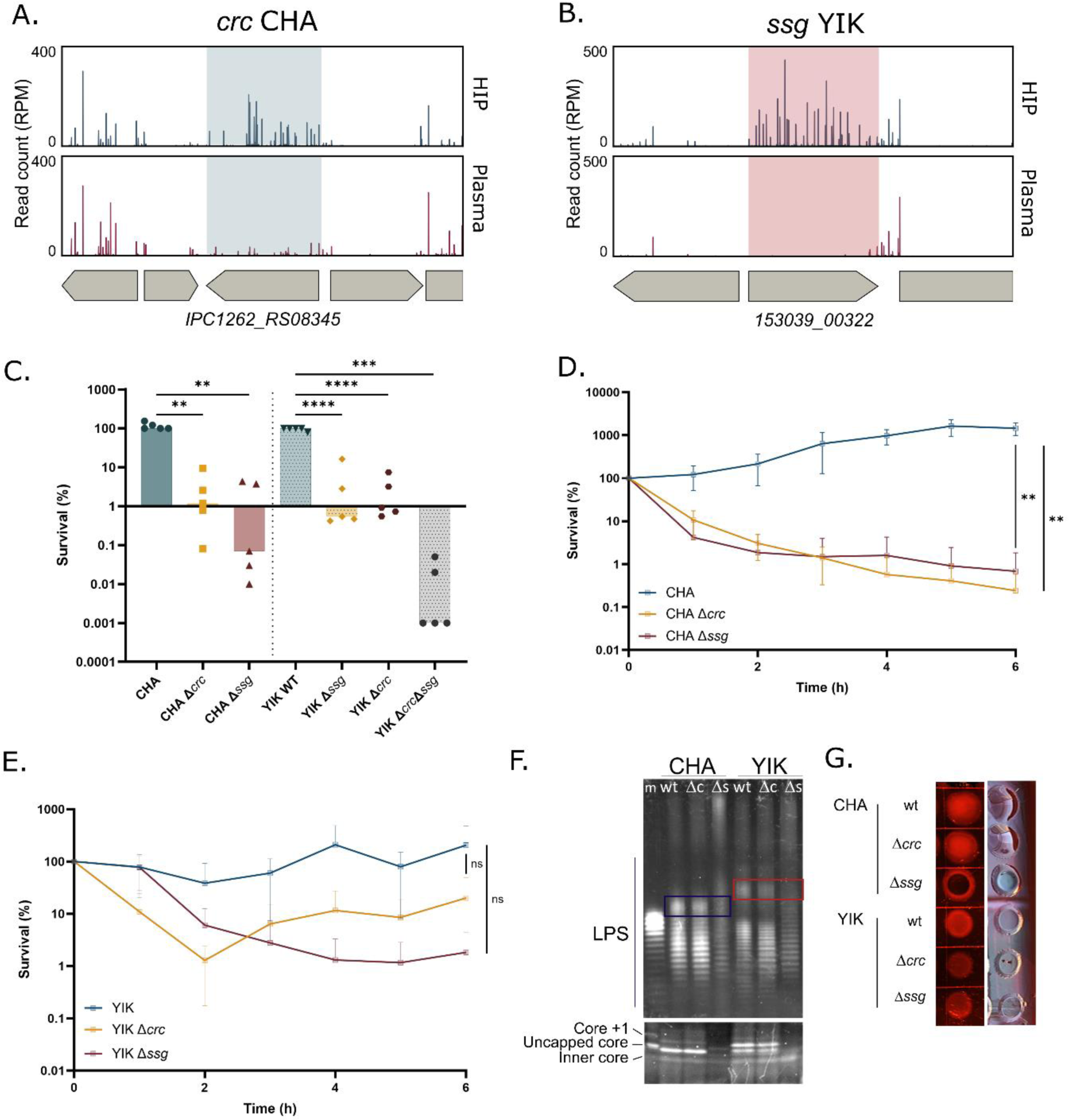
*crc* and *ssg* mutants persist in plasma. **A.** Tn-seq profile of *crc* (*IPC1262_RS08345*) genomic environment in CHA showing number of normalized reads in input (HIP) and output (plasma). **B.** Tn-seq profile of *ssg* (*153039_00322*) genomic environment in YIK showing number of normalized reads in input (HIP) and output (plasma). **C.** Survival after 3h-incubation in human pooled plasma of CHA, CHAΔ*ssg and* CHAΔ*crc and* YIK, YIKΔ*ssg*, YIKΔ*crc and* YIKΔ*crc*Δ*ssg.* Data was log-transformed for statistical analysis by a one-way ANOVA followed by a Welch and Brown-Forsythe test. **D.** Survival kinetics of CHA, CHAΔ*ssg and* CHAΔ*crc* over 6h incubation in human pooled plasma. **E.** Survival kinetics of YIK, YIKΔ*ssg and* YIKΔ*crc* over 6h incubation in human pooled plasma. **D-E.** AUC of each biological replicate was calculated, and a one-way ANOVA was applied followed by Tukey’s test. *: *p_val_*<0.05, **: *p_val_* <0.01, ***: *p_val_* <0.001, ****: *p_val_* <0.0001. ns: non-significant. **F.** LPS extraction and silver stain of CHA, CHAΔ*ssg and* CHAΔ*crc and* YIK, YIKΔssg *and* YIKΔ*crc.* The blue box shows long O-antigen in CHA and CHAΔ*crc,* absent from CHAΔ*ssg*. The red box shows a very long O-antigen in YIK and YIKΔ*crc,* absent from YIKΔ*ssg.* **G.** Congo red binding of colonies of CHA, CHAΔ*ssg and* CHAΔ*crc and* YIK, YIKΔssg *and* YIKΔ*crc*.

We therefore engineered *ssg* and *crc* deletions in both strains. The four mutants displayed survival rates between 1.6 and 4.1% (*Figure 6C*). Surprisingly, while deletions in *wzz* and *wzy* led to full bacterial elimination, *crc* and *ssg* mutants became more sensitive, but were never fully eliminated after a 3h-challenge in plasma (*Figure 6C*). We therefore investigated the killing kinetics of *ssg* and *crc* mutants. The killing curves obtained by a challenge of the different mutants in plasma for up to 6h were reminiscent of those obtained for persistent strains, PA14 or IHMA87 (9), with rapid decrease in survival within initial 2h and then a stabilization to the levels of 1-10% (*Figure 6D-E*). The plasma sensitivity after 3h of a double mutant in YIK was lower than either single mutant individually, suggesting distinct roles of *ssg* and *crc* in plasma survival (*Figure 6C*).

### Characterization of *ssg* and *crc* mutants for LPS and EPS

Literature mining suggested a possible link between *ssg* and *crc,* OSA and EPS. A *ssg* mutant in *Pseudomonas alkylphenolia* was found to lack OSA and displayed altered composition of EPS lacking fucose and mannose (62). In *P. aeruginosa*, the *ssg* mutant in PAO1 strain was impaired in auto-aggregation due to its LPS hydrophobic profile and loss of OSA while in PA14, a *ssg* mutant displayed increased susceptibility to polymyxin B and increased permeability (63–65). *Crc* encodes a post-transcriptional global regulator of carbon metabolism (66). Among other targets, Crc regulates genes involved in LPS biosynthesis (66). A *crc* deletion mutant in PAO1 had an increased proportion of LPS molecules with longer OSA compared to the parental strain.

Given the known importance of LPS in bacterial sensitivity to plasma, we first analyzed LPS content of Δ*crc* and Δ*ssg* mutants in CHA and YIK genetic backgrounds. The LPS quantity and pattern were indistinguishable between Δ*crc* mutants and their respective parental strains (*Figure 6F*), in contrast with the previous report (66). LPS extraction and staining do not enable clear discrimination between high molecular weight OSA and CPA. However, in agreement with the literature, fewer bands were observed in Δ*ssg* compared to both the WT and the Δ*crc* strains, suggesting the absence of OSA in Δ*ssg* (*Figure 6F*, (63–65)). Additionally, the Δ*ssg* mutants lacked long OSA in CHA (*Figure 6F*, blue box) and very long OSA in YIK (*Figure 6F*, red box), and exhibited decreased amounts and shift in the free LPS core, particularly striking in the CHA genetic background (*Figure 6F*). In plasma challenge experiments, Δ*ssg* formed persisters, in contrast to Δ*wzz and* Δ*wzy* mutants (*Figure 6D-E*, reviewed in (8)).

We then investigated the role of *crc* and *ssg* in EPS production using Congo red plates. CHAΔ*crc* and YIKΔ*crc* had apparent Psl production similar to that of the parental strain (*Figure 6G*). Interestingly, the colony of CHAΔ*ssg* mutant was dark red and not shiny, indicating increased Psl production and reduced alginate production (67).

We conclude that, although *crc* and *ssg* appeared as good targets in two resistant strains, their deletion led to persistence without complete bacterial elimination.

## Discussion

In this work, we used Tn-seq in three genetically and phenotypically diverse strains to investigate the convergence of molecular mechanisms interfering with the complement system cascade and subsequent bacterial killing. The data analysis highlighted no strict redundancy in plasma resistance-associated factors. This was previously reported for *K. pneumoniae* strains in serum (37). However, we could identify a few common strategies used by the different strains. In addition, results confirmed for one strain may be useful to understand plasma resistance in other strains as we highlighted for *crc* and *ssg* phenotypic analysis.

*P. aeruginosa* genomic complexity has been highlighted by various studies (22, 23) serving as resources to adapt to different environments and stresses. In addition to the lack of conservation of some potential target among strains, each strain has its own expression profile of virulence genes, which could also account for part of the diversity in molecular determinants for plasma resistance.

In plasma persistent strains, PA14 and IHMA87, the amount of transposon insertions positively affecting global survival was more important, compared to the number of hits negatively affecting the resistance of CHA and YIK. Our results suggest a highly strain-specific defense mechanism against the complement system, involving a myriad of bacterial effectors that constitute a specific cocktail of bacterial factors, shaping a strain’s resilience to complement-mediated killing (*Figure 7*).

**Figure 7.**
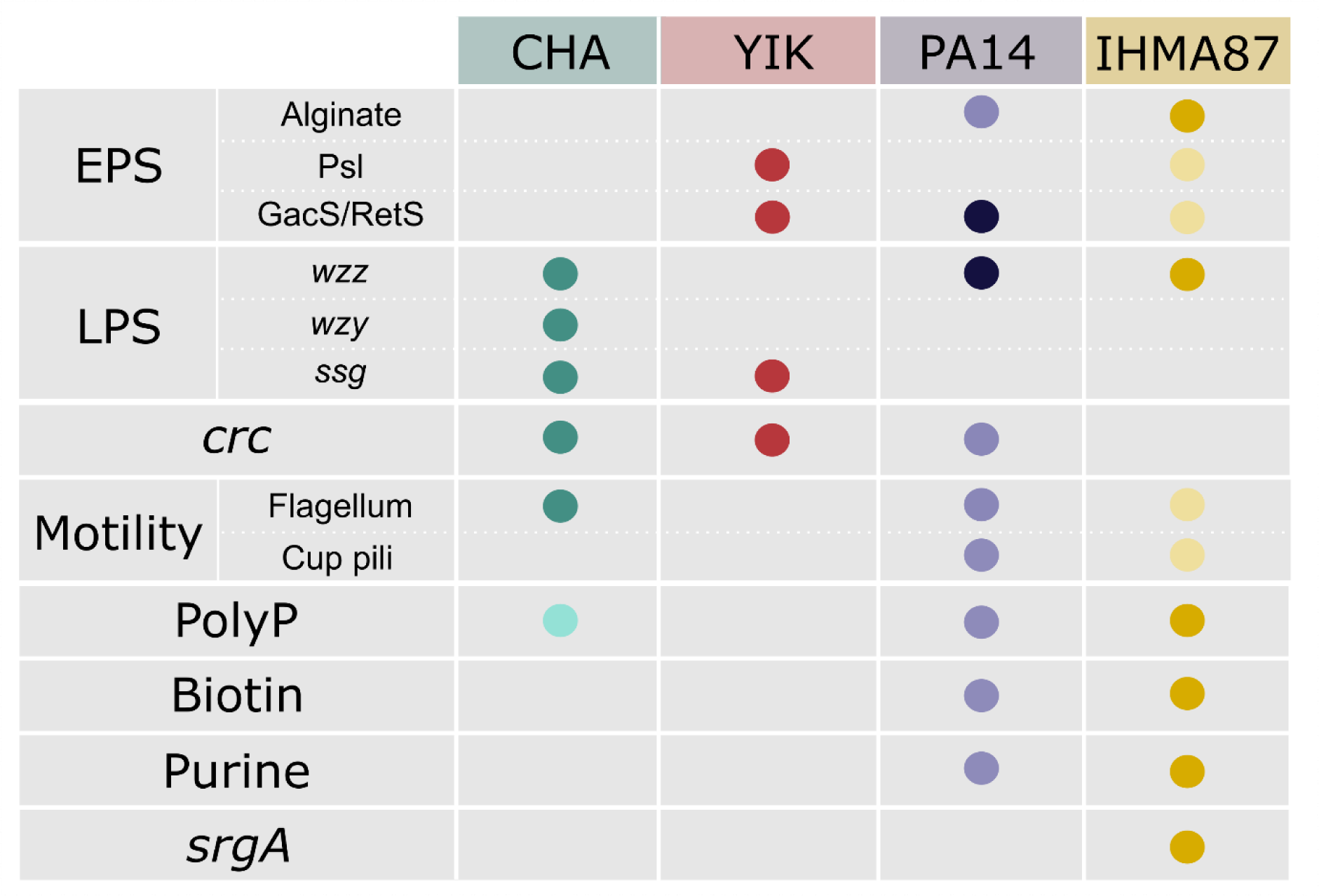
The plasma resistance of each *P. aeruginosa* strain depends on a unique combination of molecular factors. Dots represent the implication of the corresponding gene or pathways in resistance to plasma in the indicated strain either confirmed experimentally on individual mutants (darker color) or in Tn-seq data (lighter dots for CHA, PA14 and IHMA87) in this work or in (21).

Poulsen and colleagues investigated the core essential genome of nine strains of *P. aeruginosa* in five infection-relevant media including heat inactivated serum, sputum and urine (68). They propose that four strains are sufficient to establish a reliable essential genome. In our case, the variability was such that four strains were insufficient to assess the full complexity of the host-pathogen interface in plasma of *P. aeruginosa*. This raises the question of drawing conclusions from single-strain studies in host-interaction studies.

Complement system-mediated killing is initiated onto the bacterial surface and results in the formation of a pore in the outer membrane, eventually leading to bacterial lysis and death. As suggested in *Borrelia burgdorferi*, the presence of various outer membrane proteins was responsible for bacterial resistance to MAC-dependent killing (69). This was also evidenced in *K. pneumoniae*, in which the deletion of the gene coding for the outer membrane lipoprotein LppA, led to an increased bacterial susceptibility to complement-mediated killing (70). Although the exact subsequent mechanism is still under debate, two main hypotheses are drawn. The first would imply that the formation of the MAC pore on the outer membrane allows the passage of C9, subsequently forming pores in the inner membrane and causing bacterial killing. The other possible molecular mechanism implies that the formation of pore in the outer membrane destabilizes the envelope leading to the loss of envelope integrity and subsequent to bacterial death. Both hypotheses were discussed by (71). The main resistance mechanism involves the inhibition of opsonization on the bacterial envelope and degradation of complement components (70). To reduce opsonization, bacteria can secrete EPS or modify their outer membrane composition to reduce the quantity of opsonin acceptors such as OprF (36). For this reason, we expected an enrichment of outer membrane proteins as the top hits of the screen. Surprisingly, the outer membrane proteins accounted for only 9.3, 0, 2.9 and 3.2 % for CHA, YIK, IHMA87 and PA14, respectively (*Figure S6*). Conversely, all the strains had a high proportion of inner membrane-associated genes in the list of significant hits of about 25-30%. These results are in agreement with the work of Heesterbeek et al., (71) which suggests the permeabilization of the inner membrane to be the rate-limiting step in MAC-dependent killing. In addition, the screen reports many cytoplasmic hits, which may exert an indirect function onto bacterial envelope or metabolism to help the bacterium to cope with the complement system-derived envelope stress.

Global strategies, highlighted in other species such as *K. pneumoniae* or *Acinetobacter baumannii* (37, 72), were found to be redundant between strains including the production of protective exopolysaccharides and the synthesis of OSA. However, the production of motility appendages, that could protect bacteria from the complement-mediated killing effect, potentially sequestering harmful serum proteins, was never highlighted in genome-wide screens. In contrast, other screens identified the maintenance of lipid asymmetry (*mla*) as a central pathway in serum resistance (72). Mutants in the *mla* locus were also previously identified in *P. aeruginosa* PAO1 strains to be more sensitive to serum killing (73). However, in our screen, the survival of mutants in the MLA pathway was found negatively impacted by insertions in *mlaE* and *mlaF* only in the strain YIK (*Table S2*, *mlaE* - Log_2_(FC)=-1.43, *mlaF* - Log_2_(FC)=-1.21).

In this work, we also confirmed that persistence in plasma is a widespread mechanism of *P. aeruginosa* evasion, as resistant strains may become persistent by mutations in their genome. These mutations affecting bacterial surface (e.g. *ssg* mutant) or global metabolism (e.g. *crc* mutant) led to the persistence of a bacterial subpopulation of otherwise plasma resistant strains.

It is worth noting that the complement system is the main immune component responsible for the elimination of *P. aeruginosa* in the blood (9), however, it works in concert with the cellular arm of innate immunity to eliminate invaders. Therefore, genome-wide approaches involving immune cells should be considered for further studies. However, as blood from different donors cannot be pooled, this would require a high number of samples to average the immune variability between individuals. Alternatively, purified innate immune cells together with plasma could be used to screen transposon mutant’s libraries.

Overall, this work identifies many strain-specific bacterial factors as important in plasma resistance, which constitutes a valuable source of information for future work. This study also gives insight into a more global comprehension of *P. aeruginosa*/complement system interplay but also raises some concerns on the targeted antimicrobial or anti-virulence strategy, calling for more holistic studies.

## Material and methods

### Bacterial strains and genetic manipulations

Bacterial cultures were performed in Lysogeny Broth (LB) at 37°C with shaking. LB plates containing irgasan (25 µg/mL) were used to select for *P. aeruginosa*. Antibiotic concentrations used were: 75 µg/mL gentamicin for *P. aeruginosa* and 50 µg/mL gentamicin, 100 µg/mL ampicillin for *E. coli*. All bacterial strains, plasmids and primers are listed in *Tables S5-6*. Deletion mutants were engineered by amplifying approximately 500-700 bp upstream and downstream flanking regions of the gene of interest, according to (74), from genomic DNA by PCR using appropriate primer pairs (sF1/sR1, sF2/sR2). Subsequently, overlapping fragments were cloned into *Sma*I-digested pEXG2 using Sequence- and Ligation-Independent Cloning (SLIC, (75)). Using triparental mating, allelic exchange vectors were introduced into *P. aeruginosa*, with pRK600 as a helper plasmid. Selection of merodiploid clones resulting from homologous recombination was performed on LB agar plates containing irgasan and gentamicin. Merodiploids colonies were streaked onto NaCl-free LB plates with 10% sucrose (w/v) selecting for plasmid loss. Resulting clones were screened for gene deletion and antibiotic sensitivity by PCR with appropriate primer pairs.

### Generation of transposon libraries

Libraries were generated as previously described in (21). Briefly, *P. aeruginosa* strains (CHA, YIK, PA14) were grown overnight from an isolated colony at 37°C with shaking and mixed with *E. coli* SM10 carrying pBTK24 ((33), grown overnight with ampicillin). Biparental conjugation was performed, mixing equal volumes of *P. aeruginosa* and *E. coli* strains. Bacteria were pelleted by centrifugation and resuspended in fresh LB, resulting in a 5-fold concentrated bacterial suspension. Approximately 15 40µL-puddles were spotted onto dry LB plates and incubated at 37°C for 4h. Bacteria were scraped off the plates and resuspended into LB. Bacterial suspension was then spread onto irgasan- and gentamicin-containing plates and grown at 37°C. Transposon mutant colonies were collected by plate flushing with LB and scraped off using a loop. Glycerol was added to a 20% final concentration. Resulting transposon libraries were aliquoted and stored at -80°C until use.

### Plasma pools preparation

Heparinized human blood from healthy donors was provided by the French national blood institute (EFS, Grenoble, France). Fresh blood samples were centrifuged for 10 min at 1,000xg at room temperature. Plasma-containing supernatants of ten donors were pooled, filtered through a 0.45 µm membrane. Pooled plasma was then aliquoted and stored at - 80°C until needed. Prior to use, pooled plasma was thawed at 4°C and centrifuged for 10 min at 10,000 rpm on a microcentrifuge and the supernatant was filtered through a 0.22 µm membrane. To prepare heat-inactivated plasma (HIP), pooled plasma was thawed at 4°C and inactivated by a 30 min incubation at 56°C followed by a 10 min centrifugation step at 10,000 rpm and filtration through a 0.22 µm membrane.

### Screen in plasma

Screen in plasma was performed as described in (21). Briefly, transposon mutant libraries were thawed on ice and diluted in 30mL of LB. Bacteria were grown for 16h at 30°C. Bacterial culture was diluted to OD_600nm_∼0.1 and grown at 37°C under agitation. When cultures reached an OD_600nm_∼1, bacteria were harvested, resuspended in PBS supplemented with calcium and magnesium ions (PBS +/+, Thermofisher Scientific). Plasma and HIP were inoculated with a final bacterial concentration of 2.25 x 10^7^ CFU/mL (90% final plasma concentration). Initial count was determined by CFU counting and samples were incubated for 3h at 37°C on a rotating wheel. Bacterial survival at the end of the incubation was assayed by CFU counting. PA14 was spread onto LB plates to isolate transposon mutants with increased survival in plasma. After the challenge, bacteria were recovered in LB and allowed to grow overnight. Clonal populations of the corresponding wild-type strains were challenged in parallel for survival comparison. About 10^9^ bacteria were harvested for genomic DNA extraction and library constructions. Experimental procedure was performed with biological duplicates.

### Library construction, sequencing and data analysis

*Genomic DNA extraction.* Bacterial pellets were harvested from the screen and lysed by SDS-NaCl lysis (2% SDS, 0.1M EDTA pH8, 0.6M sodium perchlorate, 0.15M NaCl), followed by two successive steps of phenol-chloroform extractions. Genomic DNA (gDNA) was precipitated in 100% cold ethanol and diluted in the TE buffer (10mM Tris-HCl pH8, 1mM EDTA). gDNA was mechanically sheared at 4°C using a QSonica sonicator (Q700, 40% amplitude) for 20 min with 15 sec pulses. gDNA fragments, with a size of 150-400 bp, were then concentrated in monarch columns (PCR & DNA cleanup kit, NEB) and controlled on 2% agarose gel.

*Library construction and sequencing.* Fragmented gDNA was end-repaired using End-repair module (NEB#E6050) followed by a dA-tailing with Klenow (NEB#6053). Annealing of short and long adaptors (*Tables S6*) was performed in 10 mM MgCl_2_ in a thermocycler with a program allowing decrease of 1°C/cycle, from 95°C to 20°C. Annealed gDNA fragments were then ligated at 16°C with T4 DNA ligase (NEB#M0202). Fragments of 200 to 400 bp were selected on a 2% agarose gel and purified from gel (Monarch DNA Gel Extraction Kit, NEB#T1020L). Purified DNA-adaptor fragments were amplified using the Phusion polymerase (NEB#M0530) and a specific pair of primers (PCR1 Tn-spe direct and PCR1 Adaptor comp.). A second amplification step was performed using primers carrying the P5 and indexed-P7 illumina sequences (NEB#E7335S). At each step, double stranded DNA (dsDNA) concentration was assessed using the Qubit dsDNA HS assay kit (Q33230). Finally, the libraries quality was assessed on the Agilent Bioanalyzer 2100 (HS DNA Chips Ref #5067-4626) and sequencing was performed on an Illumina Nextseq High (I2BC, Saclay Paris).

*Data analysis.* Following quality control of the data, sequencing reads were trimmed and aligned to the corresponding genome using Bowtie (76). Readcount per feature was calculated by Htseq-count (77) and differential representation of the insertion mutants between the heat-inactivated control and the plasma sample was performed by DESeq2 (78). Concerning the analysis of Tn insertion in the intergenic regions, annotation files were created for each strain attributing the intergenic region to the downstream gene, independently of its orientation. Alignment files produced by Bowtie were then used with these new annotation files and analyzed by Htseq-count and DESeq2. All analyses were performed on the galaxy platform (38). In order to compare data from each strain, a BLAST reciprocal best (79, 80) hit analysis was performed, blasting all the proteins from each strain against PAO1 proteins with a 90% coverage and 80% identity. Reference genomes were downloaded from NCBI; EC2A-CHA: GCF_003698505.1, YIK: JAPTST000000000, UCBPP-PA14: GCF_000014625.1.

### Gene equivalent between strains

Gene equivalents among the strains of interest were determined using Blast Reciprocal Best Hits on Galaxy version 0.3.0 (usegalaxy.eu). Genes were considered equivalent when coverage was higher than 90 % and identity higher than 80 %.

### Plasma killing assay

Plasma killing assays were performed as described in (9). Bacterial cultures were inoculated with an OD_600nm_ of 0.1 and grown at 37°C with shaking until it reached an OD_600nm_ of 1. Bacteria were harvested and resuspended in PBS +/+ and plasma was inoculated with a final bacterial concentration of 2.25 x 10^7^ CFU/mL (90% final plasma concentration). Initial and final bacterial loads were assessed by CFU counting at indicated times.

### LPS Extraction and visualization

LPS was extracted by the hot phenol-water method as described previously (81, 82) with some modifications. Briefly, 30 mL of bacterial culture were pelleted by centrifugation and resuspended in 1mL of 20mM Tris pH8, 20mM MgCl_2_, 50mM NaCl buffer. Sonication was performed for 20 min at 4°C (10 sec on/10 sec off, 40% amplitude) using a QSonica Oasis-180. In order to eliminate proteins and nucleic acids, treatment with DNase (Euromedex 1307, 50µg/ml) and RNase (Sigma R6513, 40µg/ml) was carried out for 1h 37°C followed by Proteinase K (Roche-311879001) treatment for 1h at 65°C and an overnight incubation at 37°C. Then, two successive hot phenol-water extractions (Invitrogen 15594-04) were carried out. Briefly, equal volume was added, vigorously mixed and incubated 10 min at 65°C, then centrifuged 10 min 20,000 xg at 4°C. The aqueous phase was precipitated by 0.5M sodium acetate (pH 5.2) with 10 volumes of cold ethanol, and stored at -20°C. After centrifugation 10 min 20,000 xg at 4°C, pellets were washed with water and dialyzed. Final purified LPS product was lyophilized and stored at -20°C. Freeze-dried samples were resuspended in SDS-PAGE buffer and boiled for 5 min. Samples were loaded onto a NuPAGE™ Bis-Tris 4-12 % gel (Invitrogen). LPS were stained using the ProQ Emerald 300 lipopolysaccharide gel stain kit (Molecular Probe P20495).

### CONGO Red Plates

Five microliters of culture at OD_600nm_ of 1 were spotted onto BHI agar plate (Brain Heart Infusion Broth - Sigma 53286) containing 5% sucrose and 0.8 g/L Congo red (Sigma C6767). Plates were incubated for 24h at 37°C and at 4°C before imaging.

### Statistical analysis

Statistical analyses were performed using Sigmaplot and Graphpad prism 9 software. The test used are specified in figure legends. Survival data were log_10_-transformed. Figures were performed using Graphpad Prism 9 and Inkscape.

## Data availability

The Tn-seq data generated in this work is available at GEO under the series accession numbers GSE287536.

## Acknowledgement

The work was supported by grants from Agence Nationale de la Recherche (ANR-15-783 CE11-0018-01), the Laboratory of Excellence GRAL, financed within the University Grenoble Alpes graduate school (Écoles Universitaires de Recherche) CBH-EUR-GS 785 (ANR-17-EURE-0003), the Fondation pour la Recherche Médicale (Team FRM 2017, 786 DEQ20170336705) to I.A. The sequencing part has benefited from the facilities and expertise of the high throughput sequencing core facility of I2BC (Centre de Recherche de Gif – http://www.i2bc.paris-saclay.fr/). M. JM received a Ph.D. fellowship from the French Ministry of Education and Research. The authors thank Peter Panchev and Emily Rey for their help for transposon mutant libraries construction, and Joanna Goldberg and Dina A. Moustafa for their valuable discussion.

